# Target-responsive vasoactive probes for ultrasensitive molecular imaging

**DOI:** 10.1101/786079

**Authors:** Robert Ohlendorf, Agata Wiśniowska, Mitul Desai, Ali Barandov, Adrian L. Slusarczyk, Alan Jasanoff

**Affiliations:** Department of Biological Engineering, Massachusetts Institute of Technology, 77 Massachusetts Ave., Rm. 16-561, Cambridge, MA 02139; Harvard-MIT Health Sciences & Technology, Massachusetts Institute of Technology, 77 Massachusetts Ave., Rm. 16-561, Cambridge, MA 02139; Department of Brain & Cognitive Sciences, Massachusetts Institute of Technology, 77 Massachusetts Ave., Rm. 16-561, Cambridge, MA 02139; Department of Nuclear Science & Engineering, Massachusetts Institute of Technology, 77 Massachusetts Ave., Rm. 16-561, Cambridge, MA 02139

## Abstract

The ability to monitor molecules volumetrically throughout the body could provide valuable biomarkers for studies of healthy function and disease, but noninvasive detection of molecular targets in living subjects often suffers from poor sensitivity or selectivity. Here we describe a family of potent imaging probes that can be activated by molecules of interest in deep tissue, providing a basis for mapping nanomolar-scale analytes without the radiation or heavy metal content associated with traditional molecular imaging agents. The probes are reversibly-caged vasodilators that induce responses detectable by hemodynamic imaging; they are constructed by combining vasoactive peptides with synthetic chemical appendages and protein blocking domains. We use this architecture to create ultrasensitive biotin-responsive imaging agents, which we apply for wide-field mapping of targets in rat brains using functional magnetic resonance imaging. We also adapt the sensor design for detecting the neurotransmitter dopamine, illustrating versatility of this approach for addressing biologically important molecules.

## INTRODUCTION

Mapping molecular species with minimal invasiveness in whole living organisms could provide insight into diverse physiological functions and their disruption in disease. Fluorescent sensors are commonly used to monitor molecular processes at high resolution *in vivo*^1,2^, but detection of fluorescent probes in most organisms is limited by optical scattering and absorption to superficial and often decontextualized regions of the body^3–6^. Sensitive detection of probes at subnanomolar concentrations in deep tissue is possible using nuclear imaging, but analyte-responsive probes for nuclear techniques are not available, so these methods can only measure localization and kinetics of the tracers themselves^7^. Magnetic resonance imaging (MRI) contrast agents are detectable in deep tissue, and can be sensitized to a variety of biologically-relevant targets^8^, but imaging agents for MRI are usually only detectable at micromolar concentrations, or at similar mass doses when nanoparticle contrast agents are used. This creates challenges for delivery of the probes, increases the potential for physiological disruptions or toxicity, and often means that only high analyte levels can be detected.

Vasoactive probes (vasoprobes) are a new class of imaging agent that can be detected at an organ- or organism-wide scale using diverse noninvasive imaging modalities, without the need for radioactive or metallic components^9^. Effective vasoprobes can be derived from potent vasodilatory peptides that induce relaxation of the vascular smooth muscle cells (VSMCs) that surround most blood vessels. This in turn leads to spatiotemporally localized changes in blood flow, volume, and oxygenation that produce contrast changes in MRI, ultrasound, nuclear imaging, and optical imaging, analogous to the widely-exploited hemodynamic bases of functional neuroimaging methods^10–13^. Recent results show that vasoprobes can be detected at nanomolar concentrations in the rodent brain and that both secretion and proteolytic uncaging of vasoprobes give rise to specific signatures in imaging^9^. Vasoprobes thus combine the versatility of paramagnetic MRI contrast agents with sensitivity over 1,000 times greater^8^. We therefore reasoned that the vasoprobe contrast mechanism could be harnessed to create potent sensors for biologically-important molecules, and that target-responsive vasoprobes could facilitate detection of molecular species at the nanomolar concentrations characteristic of many signaling molecules, biochemical markers, and therapeutic agents.

In this study, we present a vasoprobe-based strategy for sensing biologically important molecules by hemodynamic imaging *in vivo*. We engineer activatable vasoprobes for analyte targeting (AVATars) with the aid of a cell-based *in vitro* bioassay, and show that the resulting imaging probes permit target-responsive detection of nanomolar-scale molecular species in live rat brains. We use our first AVATar to map biotin derivatives *in vivo*, demonstrating non-invasive detection of dilute biotinylated epitopes in intact tissue. We then demonstrate the generality of our design by creating a second AVATar for detection of the important neurotransmitter dopamine, indicating the potential of vasoprobe technology for addressing a wide variety of molecular phenomena in the brain and other organs.

## RESULTS

### Platform for AVATar construction

We chose to derive AVATars from the potent vasodilator pituitary adenylate cyclase-activating polypeptide (PACAP), a 38-residue peptide that activates G protein-coupled receptor-dependent cyclic adenosine monophosphate (cAMP) signaling in VSMCs with half-maximal effective doses (EC_50_) in the sub-nanomolar range^14^ (**Figure 1a**). We envisioned an AVATar design wherein a PACAP derivative is labeled with a tethered version of the target molecule of interest, which in turn interacts with a protein domain that selectively binds either the tethered target or the target molecule itself, in competition (**Figure 1b**). In the absence of the target molecule, the protein domain would remain bound to the PACAP moiety via the tethered ligand, blocking the PACAP moiety from activating its receptors. The presence of elevated concentrations of the target molecule in tissue, however, would cause release of the PACAP derivative from the blocking protein, activating the AVATar and evoking a local hemodynamic imaging signal.

**Figure 1.**
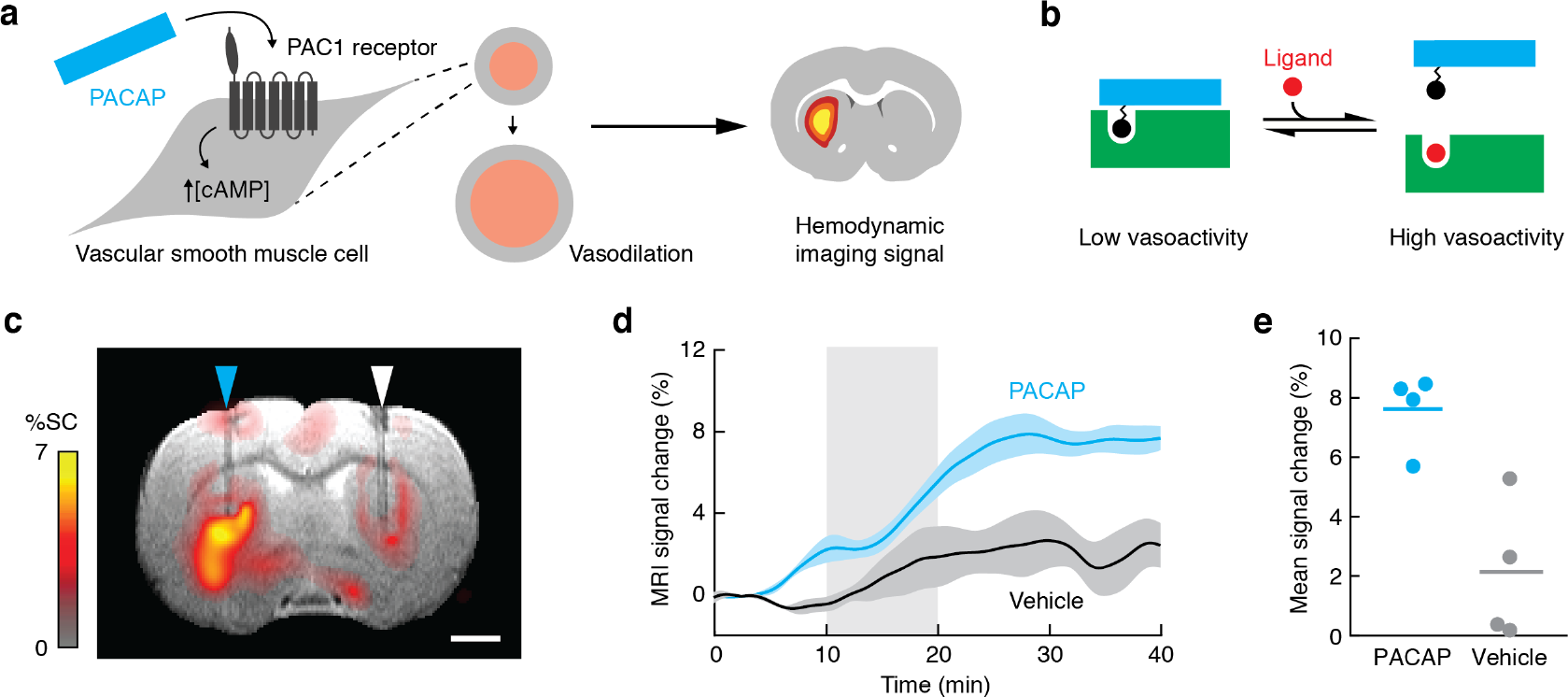
Platform for AVATar construction. **(a)** PACAP functions as a vasoprobe by binding to receptors on vascular smooth muscle cells and activating a cAMP-dependent signaling cascade, causing blood vessel dilation and a hemodynamic imaging signal. **(b)** AVATar architecture: A tethered version of the target molecule is attached to PACAP peptide (blue); binding of a blocking domain (green) to the tethered target prevents receptor activation. Elevated free target molecule concentrations free active PACAP from the blocking interaction. **(c)** Map of mean MRI signal change following intracranial delivery of 1 μM PACAP (cyan arrowhead) compared to aCSF vehicle (white arrowhead), overlaid on an anatomical scan. Scale bar = 2 mm. **(d)** Mean MRI time courses during delivery of 1 μM PACAP (blue) or aCSF vehicle (black). Shading represents SEM (*n* = 4). **(e)** Average MRI signal change 30-40 minutes after PACAP or vehicle injection (horizontal lines = mean values).

To verify that PACAP could be a suitable basis for constructing such sensors, we first examined whether this peptide could function as an effective vasoprobe in the living rat brain (**Figure 1c**). Intracranial delivery of 1 μM PACAP caused a mean hemodynamic blood oxygenation level-dependent (BOLD) signal change of 7.6 ± 0.6% in *T*_2_* relaxation-weighted MRI compared to only 2.1 ± 1.2% in response to control injections of artificial cerebrospinal fluid (aCSF) vehicle (**Figure 1d-e**); this difference was highly significant with *t*-test *p* = 0.007 (*n* = 4). PACAP-dependent responses could also be visualized over sequential blocks of injection (**Supplementary Figure 1**). These results suggested that PACAP can indeed function as a platform for AVATar design.

### Engineering a PACAP-based AVATar

We decided to apply the AVATar principle initially for sensing the small molecule biotin. Because biotin is widely used as a labeling moiety, a vasoactive biotin sensor can be immediately useful as a means of detecting endogenously or exogenously biotin-labeled species in tissue, thus facilitating radiation-free *in vivo* analogs of powerful assays and histology methods^15^, as well as pretargeted imaging approaches^16^. With a molecular weight of 244 Da, biotin is similar in size and characteristics to many endogenous signaling molecules and metabolites, so we also anticipated that an imaging sensor for biotin could subsequently be adapted for detection of other targets. The tetrameric 53 kDa protein streptavidin (SA) and its close relatives bind tightly to both free and functionalized biotin derivatives^17^, and are obvious candidates to act as blocking domains. The AVATar design also requires site-specific tethering of biotin to PACAP in a way that does not severely interfere with receptor activation, allows for binding of the blocking domain, and results in a sufficient activity change between blocked and unblocked states, such that substantial vasodilation occurs only in the unblocked state. We used a combination of structural data^18^ and previous structure-function analysis^14,18–20^ results to identify candidate biotin attachment sites likely to satisfy these criteria (**Figure 2a**). A set of PACAP derivatives was then prepared, for which each of these sequence positions was replaced with a lysine-biotin residue.

**Figure 2.**
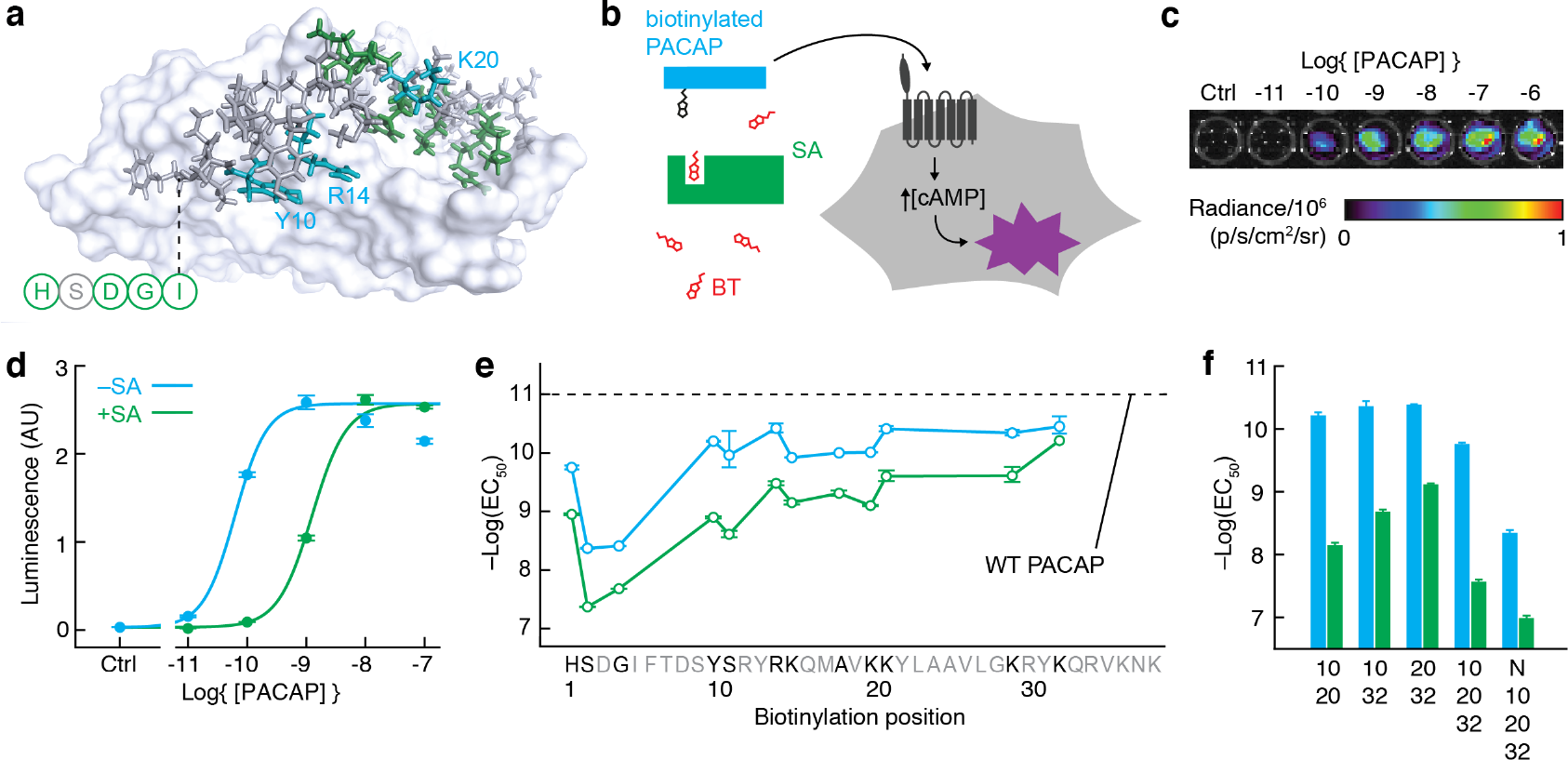
Engineering a PACAP-based AVATar. **(a)** Identification of candidate ligand attachment sites on PACAP: Structure of PACAP in complex with a PAC1 receptor fragment^18^, highlighting residues whose substitution blocks (green) or does not affect (cyan, labels) peptide bioactivity, according to previous structure-activity studies. N-terminal residues absent from the structure indicated schematically (circles). **(b)** Schematic of PAC1-based bioassay for testing biotinylated PACAP constructs (cyan), their blocking by streptavidin (green), and release by free biotin (red). Unblocked PACAP derivatives can bind the PAC1 receptor (dark gray) expressed in CHO cells, triggering cAMP production and activating a luminescence reporter (purple). **(c)** Raw luminescence recorded from the assay cells under titration with wild type PACAP. **(d)** Quantification of luminescence output and determination by curve-fitting of EC_50_ values for a biotinylated PACAP derivative (PACAP-10-BT) in the presence (green) and absence (cyan) of 200 nM streptavidin (SA). **(e)** Screening for PACAP residues that tolerate modification: EC_50_ values for receptor activation in the absence (cyan) or presence (green) of SA by PACAP-biotin conjugates as a function of sequence position (untested positions in gray). EC_50_ of wild-type (WT) PACAP indicated by the dashed line. **(f)** EC_50_ values of multiply-biotinylated PACAP conjugates modified at the positions noted (N = N-terminus), in the absence (cyan) and presence (green) of SA. Error bars in **d**-**f** denote SD of duplicate measurements.

To rapidly screen the modified peptide variants, we established a luminescence assay in mammalian cells that models PACAP-related vasoactivity by quantitatively reporting cAMP production in response to activation of the PACAP receptor PAC1 (**Figure 2b,c**). Titration of PACAP derivatives in this assay enabled EC_50_ values to be obtained for each variant, in both the presence and absence of SA (**Figure 2d**). With the exception of several residues near the N-terminus, biotinylation of most PACAP residues is well tolerated in the absence of SA, with variants displaying EC_50_ values only modestly worse than the value of 10.8 ± 0.2 pM exhibited by wild-type PACAP (**Figure 2e**). Losses of potency by ~300-fold upon biotinylation of positions 2-5 is consistent with previous studies reporting the requirement for these residues in receptor activation^14,18^. Of the PACAP derivatives evaluated, those with biotinylation of sites in the mid-region of the peptide produced the greatest SA-dependent dynamic range in receptor activation. In particular, a PACAP variant biotinylated at residue Y10 (PACAP-10-BT) provides the best combination of preserved bioactivity and sensitivity to SA binding, with EC_50_ values of 63 ± 1 pM and 1.24 ± 0.05 nM in the absence and presence of SA, respectively, corresponding to a 20-fold change in potency (**Figure 2d**). A PACAP variant with positions 10 and 20 simultaneously biotinylated (PACAP-10,20-BT) produced an even greater dynamic range, with an EC_50_ of 61 ± 7 pM that is reduced by a factor of 115 ± 22 in the presence of SA (**Figure 2f**). Addition of further PACAP biotinylation sites did not substantially improve on this sensitivity, however, nor did modifications to the biotin-peptide linker length or repurification to remove contaminants from peptide synthesis (**Supplementary Table 1**).

### Detecting biotin with nanomolar sensitivity *in vitro*

Based on these results, we designed a functional biotin-responsive sensor, BT-AVATar. The combination of PACAP-10,20-BT and SA could in principle function as a suitable sensor in itself, but the characteristic dissociation time of SA from biotin (~7 hours)^21^ is too long for meaningful applications based on the competition principle of **Figure 1b**. To achieve effective biotin sensing, we therefore replaced the tethered biotin residues in PACAP-10,20-BT with desthiobiotin, a similar compound that binds SA with 10,000-fold lower affinity^22^ than biotin and can be easily displaced by it^23^. The resulting peptide, PACAP-10,20-dBT, is not as effectively blocked by SA as PACAP-10,20-BT, but restoration of further modification sites addresses this problem (**Supplementary Figure 2**). A variant in which tethered desthiobiotins were attached at positions 10, 20, and 32 proved optimal; this PACAP-10,20,32-dBT displayed EC_50_ values of 130 ± 2 pM and 8.2 ± 0.6 nM in the absence and presence of SA, a robust 63-fold difference in potency. BT-AVATar was then formed by mixing 1 nM PACAP-10,20-32-dBT with two-fold molar excess of SA.

Titration of BT-AVATar with varying amounts of biotin in the PAC1 bioassay reveals an EC_50_ for biotin detection of 120 ± 10 nM (**Figure 3a**). This is sensitive enough to detect typical biotinylation levels associated with biotin display or labeling technologies^15^, but not so sensitive as to be triggered by endogenous plasma biotin levels^24^. The response of BT-AVATar to biotin is reversible over repeated cycles of excess SA and biotin addition (**Figure 3b**).

**Figure 3.**
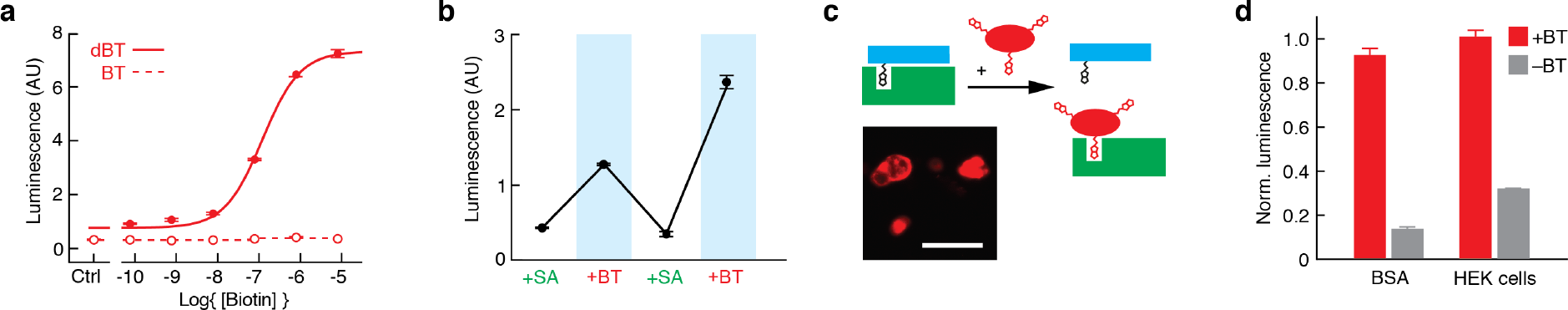
Validation of BT-AVATar *in vitro*. **(a)** Titration of BT-AVATar containing dBT-modified PACAP with biotin (solid line) showing activation with EC_50_ of 120 ± 10 nM. An equivalent variant of the sensor containing BT-modified PACAP (dashed line) is not activated by biotin. **(b)** Biotin vasosensor activity over repeated cycles of excess SA and biotin addition. **(c)** Schematic showing AVATar activation by biotinylated proteins or cells. Inlay shows biotinylated HEK cells stained with fluorescent streptavidin. Scale bar = 50 μm. **(d)** BT-AVATar is robustly activated by biotinylated bovine serum albumin and HEK cells (red) but not by unbiotinylated controls (gray). All error bars denote SD of duplicates.

The purpose of a biotin sensor is to map biotin-labeled targets *in vivo*, analogous to a variety of biochemical and histological techniques, but in intact tissue. To test functionality of BT-AVATar for this purpose, we used the PAC1 bioassay to evaluate the sensor’s *in vitro* responses to biotin moieties immobilized on molecules and cells (**Figure 3c**, **Supplementary Figure 3**). As expected, BT-AVATar is strongly activated by both biotinylated bovine serum albumin and HEK293 cells, exhibiting increases in PAC1 receptor activation by 675 ± 69% and 227 ± 30%, respectively, versus minimal responses to unbiotinylated controls (**Figure 3d**). These results show that BT-AVATar maintains sensitivity and selectivity for biotin even in biological environments, suggesting the suitability of vasoprobe-based sensors for applications *in vivo*.

### Ultrasensitive AVATar-based molecular imaging *in vivo*

To demonstrate the ability of BT-AVATar to perform sensitive molecular imaging of biotinylated targets in live animals, we implanted biotinylated and nonbiotinylated cells at symmetric sites in rat cortex and performed molecular imaging using the probe (**Figure 4a**). In order to survey a large region of the brain with minimal invasiveness, including but not limited to the cell implantation sites, BT-AVATar was infused distally into the cerebrospinal fluid (CSF) at the rostral extent of the cortex. Dissemination of imaging agents introduced using this route was verified by injecting the conventional MRI contrast agent gadolinium diethylenetriaminepentaacetic acid (Gd-DTPA) and then visualizing the results by *T*_1_-weighted imaging. Indeed, a volume of 40 μL of 200 mM Gd-DTPA spread throughout the anterior quadrant of the brain, enhancing contrast across a large volume including the cell implantation site, as well as additional regions of the olfactory bulb, frontal cortex, prelimbic cortex, motor cortex, and cingulate cortex (**Figure 4b**).

**Figure 4.**
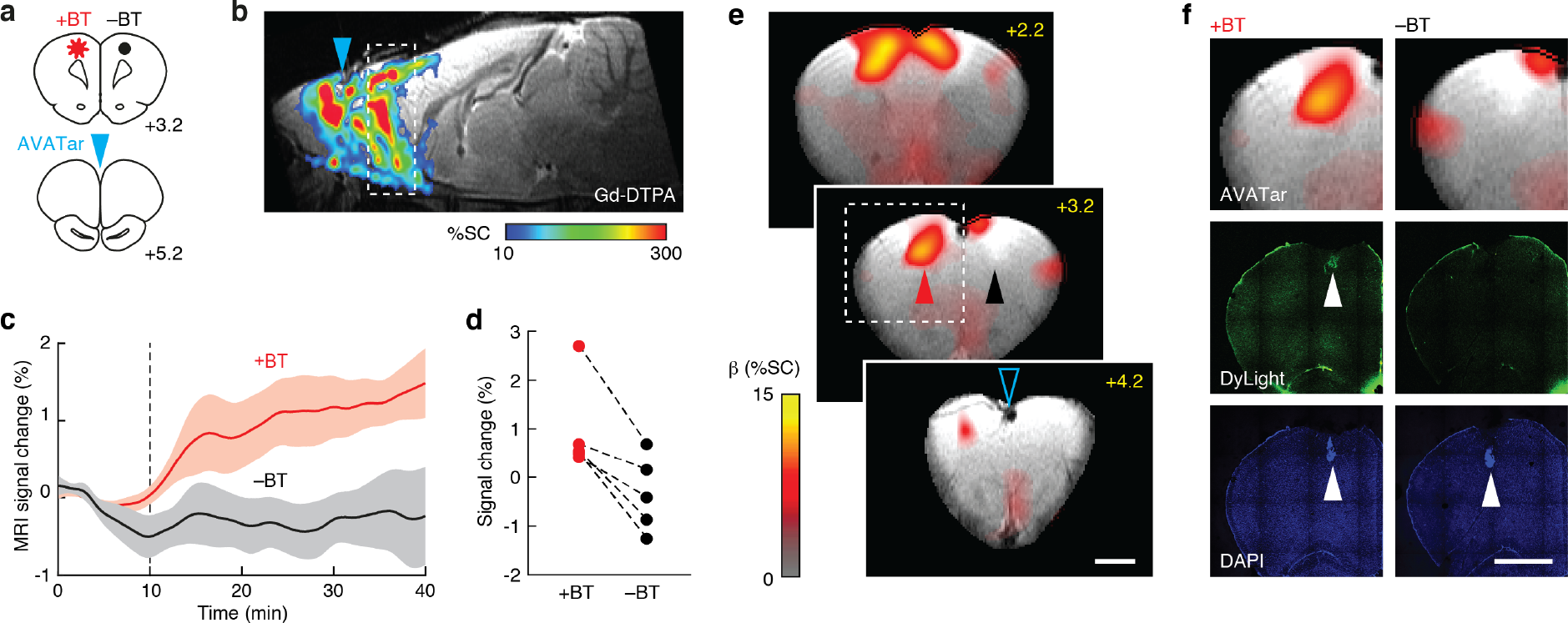
*In vivo* molecular imaging with BT-AVATar. **(a)** Brain atlas^49^ sections (bregma coordinates noted) showing positions of test (+BT) and control (−BT) cell implants and intra-CSF AVATar infusion for *in vivo* mapping experiments. **(b)** Wide-field probe delivery illustrated using Gd-DTPA infused at the AVATar infusion position (cyan arrowhead). Color scale illustrates signal changes (%SC) produced by the contrast agent. **(c)** Signal change over time near test and control cells. Dashed line indicates start of AVATar infusion. Shading denotes SEM of *n* = 5. **(d)** Signal enhancements in individual animals at +BT and −BT sites. **(e)** Map of regression coefficients (β, in units of %SC) illustrating responses near +BT cell implant locations (red arrowhead) but not −BT controls (black arrowhead). BT-AVATar was infused 1 mm anterior to the open cyan arrowhead. Displayed brain slices correspond to the dashed region in **b**. Scale bar = 2 mm. **(f)** Comparison of MRI results with histology shows overlap of BT-AVATar signal with position of test implants (left images) but not control implants (right images). Top row: closeup of dashed region in **e** (left), containing +BT cells, or symmetric region containing −BT cells (right). Middle row: selective activation of BT-AVATar by biotinylated cells is confirmed by visualization of biotinylated cells (left, arrowhead) but not unbiotinylated cells (right) following infusion of fluorescent BT-AVATar formulated with DyLight-labeled SA. Bottom row: visualization of both test and control xenografts (arrow-heads) by DAPI staining. Scale bar = 2 mm.

In the area of the implanted cells, BT-AVATar produces MRI signal changes with amplitudes that clearly distinguish biotinylated from control xenografts (**Figure 4c**). Mean hemodynamic responses average 1.0 ± 0.4% in the neighborhood of biotinylated cells, versus −0.3 ± 0.4% near control cells (**Figure 4d**), a significant difference (paired *t*-test *p* = 0.01, *n* = 5). Temporal analysis of the wide-field imaging data produces a map of corresponding changes over a wide field of view; this map indicates signal changes clustered primarily near the area of biotinylated cell implantation, despite wide-field administration of BT-AVATar (**Figure 4e**). Comparison of the molecular imaging map with histological results confirms colocalization of the AVATar-mediated signal with biotin-containing xenografts (**Figure 4f**). Estimation of biotin content in the test cell implants indicates that biotin concentrations of about 150 nM were present (**Supplementary Figure 4**), far below analyte levels detectable by conventional paramagnetic MRI contrast agents. These results thus demonstrate that nanomolar target concentrations can be imaged using BT-AVATar in conjunction with wide-field CSF delivery methods in rat brain.

### AVATar-based neurotransmitter detection

The AVATar design can be generalized for sensing a wide variety of molecular targets. Neurochemicals are of particular interest because of their distinct functional roles in the nervous system and their importance in health and disease. Although neurotransmitter-responsive probes for MRI have been reported, they are not sensitive enough to report the submicromolar extracellular concentrations characteristic of many species. To adapt the sensing mechanism of BT-AVATar for neurotransmitter detection, we mutated PACAP residues Y10 and K20 to cysteines and conjugated the resulting thiol groups to synthetic thiol-reactive derivatives of dopamine (DA), a neurochemical involved in addiction, motivated behavior, and motor control (**Figure 5a**, **Supplementary Figure 5)**. The resulting conjugate showed an EC_50_ value of 2.5 ± 0.6 nM for PAC1 receptor activation in the cell-based assay (**Figure 5b**), indicating that tolerance for modification of PACAP at the 10 and 20 positions is general. The dopamine sensor DA-AVATar was formed by mixing the modified peptides with excess immunoglobulin G directed against the tethered neurotransmitter moieties. In this sensor, the DA-specific IgG functions as a protein blocking domain according to the design of **Figure 1b**, analogously to the SA component in BT-AVATar, and with blocking reversible by addition of excess DA (**Figure 5c**). Titration of DA-AVATar with DA indicates that the sensor has an EC_50_ value of 5.6 ± 0.5 μM for its analyte, with a limit of detection below 1 μM (**Figure 5d**).

**Figure 5.**
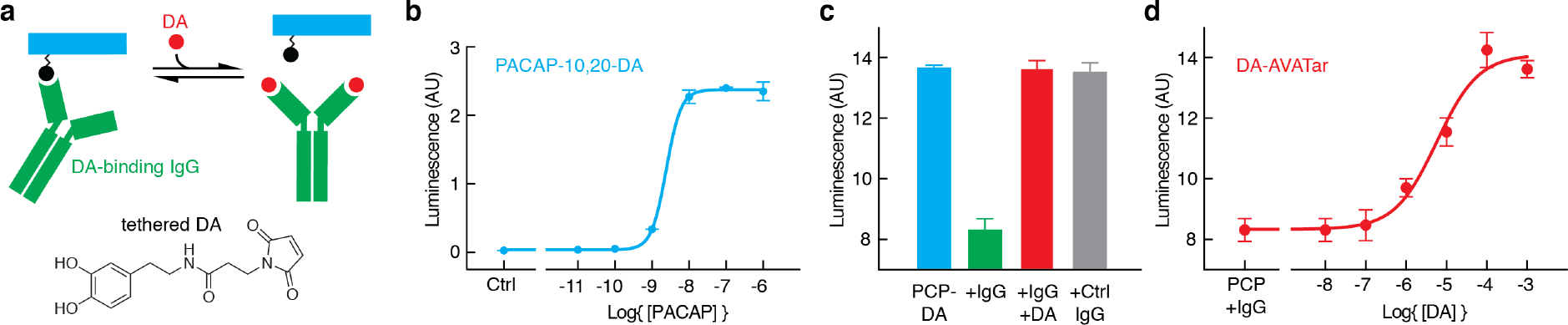
A neurotransmitter-sensitive AVATar. **(a)** Mechanism of dopamine-sensitive AVATars: PACAP is modified with a tethered dopamine (DA) analog (black), and a DA-binding IgG (green) functions as the blocking domain. Free neurotransmitter (red) competes with this interaction, releasing active PACAP (cyan). Structure of thiol-reactive tethered DA shown at bottom. **(b)** Titration of PACAP-10,20-DA in the PAC1 cell assay, showing efficient receptor activation. **(c)** PAC1 receptor activation by 1 nM PACAP-10,20-DA in the absence (cyan) or presence (green) of 600 nM DA-binding IgG. Activity is restored by addition of excess DA (red). Control IgG (gray) does not block PACAP activity. **(d)** Titration of the DA response of 1 nM DA-AVATar in the PAC1 activation assay. DA activates DA-AVATar with an EC_50_ of 5.6 ± 0.5 μM. Error bars denote SD of duplicates.

## DISCUSSION

Our work thus introduces AVATars as a versatile nonmagnetic and radiation-free imaging probe architecture for detecting molecular targets with nanomolar sensitivity in deep tissue. These probes are applied at concentrations more than 1,000-fold lower than conventional MRI agents and comparable to some nuclear imaging probes, making them particularly suitable for noninvasive imaging of the many biological species that are present only at submicromolar concentrations *in vivo*. We designed the biotin-sensitive imaging agent BT-AVATar, which is able to visualize biotin-labeled cell populations following wide-field delivery of the probe in live rat brains. In the future, BT-AVATar could also be used for direct detection of other biotinylated biomolecular and cellular targets, including targets biotinylated *in vivo*^25^, or for indirect detection in conjunction with pretargeted antibody-biotin conjugates. We demonstrate adaptability of the AVATar architecture by designing an agent that detects the neurotransmitter DA. Incorporation of alternative tethered ligands and cognate binding domains could permit a variety of further vasoprobe-based sensors to be constructed, in some cases leveraging well-characterized components of previously described fluorescent imaging probes^26–35^.

The high sensitivity of vasoprobe-based agents for their targets arises from their exploitation of physiological amplification pathways built into the vasculature. Quantification of analyte levels using this approach should be possible in contexts where calibration of AVATar-mediated signals can be performed with respect to known target concentrations. The PAC1 activation assay used here could provide an external standard for this. For instance, if we suppose that the biotin-triggered BT-AVATar responses of **Figure 4** constitute about half the maximum response of ~15% observed in similar hemodynamic imaging experiments, then we would crudely but correctly infer that the amount of biotin detected approximately matches the *in vitro* EC_50_ of BT-AVATar for biotin. This external calibration approach could be improved by performing explicit measurements of maximal hemodynamic changes *in situ*, using established techniques^36–38^.

The spatiotemporal properties of AVATar-based analyte sensing likewise follow largely from their unique hemodynamic mechanism. The probes are compatible with diverse hemodynamic imaging methods, including magnetic, nuclear, ultrasonic, and optical approaches that in some cases provide spatial detail on the order of single blood vessels (< 100 μm). The temporal resolution provided by AVATars is likely to be limited by hemodynamic response time courses on the order of seconds, and may be further limited by the kinetics of the competitive interactions between analytes and the blocking domains used in the sensor designs presented here. For neurotransmitter sensing, kinetic characteristics of AVATars may thus be comparable to previous neurotransmitter-sensitive MRI probes, which display response time constants of tens of seconds; these are favorable with respect to receptor displacement strategies used in positron emission tomography, which generally require measurements lasting tens of minutes.

The probe architecture we describe here entails the combination of ~5 kDa PACAP derivatives with proteins that range from 60 to 150 kDa in molecular weight. Even the largest of these has a hydrodynamic radius smaller than most nanoparticle-based imaging agents. The relatively small size and high potency of AVATars should facilitate delivery of these probes to a number of tissue types in which they are predicted to elicit analyte-dependent hemodynamic responses^39–44^. The brain is of particular interest because of its important chemical signaling systems and inaccessibility to most diagnostic agents. Interestingly, PACAP derivatives have been shown in earlier studies to spontaneously permeate the blood-brain barrier, suggesting the possibility of completely noninvasive delivery of PACAP-based sensors to the central nervous system. Although this avenue remains to be explored, the experiments presented here already demonstrate that AVATars can access large fields of view in the rodent brain following mini-mally-invasive intra-CSF infusion. Similar probes applied using such delivery approaches could thus provide unprecedented capability for monitoring molecular targets throughout much of the body.

## METHODS

### Plasmids

Lentiviral plasmids were constructed using the Golden Gate method^45^ in the pLentiX1 Zeo plasmid (Addgene #17299) with its kanamycin resistance replaced by the ampicillin resistance cassette. The Glo22F gene encoding an cAMP-modulated luciferase (Promega, Madison, WI)^46^ was cloned followed by an internal ribosome entry site, a Cerulean fluorescent protein, a 2A viral sequence^47^ and a puromycin resistance marker. The PAC1 gene was cloned followed by an internal ribosome entry site, an mKate fluorescent protein, a 2A viral sequence and a blasticidin resistance marker. Lentiviral helper plasmids pMD2.G (Addgene #12259, Cambridge, MA) and psPAX2 (Addgene #12260) were gifts from Didier Trono.

### Mammalian cell culture

CHO K1 cells were purchased from Sigma-Aldrich (St. Louis, MO) and cultured in 90% F10 medium supplemented with 10% fetal bovine serum (FBS), 100 units/mL penicillin, and 100 μg/mL streptomycin. Cells were frozen in freezing medium composed of 80% F10 medium, 10% FBS and 10% dimethylsulfoxide.

### Lentivirus production and cell line generation

293FT cells (Thermo Fisher Scientific, Waltham, MA) were seeded into 6-well plates at 1 million cells per well and transfected using Lipofectamine 2000 (Life Technologies, Grand Island, NY) according to instructions at sub-confluence. Co-transfection of 0.5 mg pMD2.G, 1 mg psPAX2 and 1 mg of the lentiviral plasmid of interest was performed with 6.25 ml Lipofectamine 2000 reagent. Virus-containing supernatant was collected after 48 and 72 h, filtered through 0.45 mm filters, and used for infection. Supernatants were stored at 4 °C for up to a week. CHO K1 cells were seeded into 24-well plates at 40,000 cells per well in the presence of 4 mg/ml Polybrene in 50% fresh medium and 50% viral supernatants containing both Glo22F and PAC1 viruses. The medium was replaced with fresh viral supernatants daily for 4 days. Selection was performed using both antibiotic resistance and fluorescent markers for each lentivirus. Beginning on day 3 after initial infection, 10 mg/ml blasticidin and 1 mg/ml puromycin (Life Technologies) were added to the medium for selection and selection was continued until all cells expressed the appropriate fluorescent markers.

### PAC1 luminescence assay

Production of intracellular cAMP by the PACAP receptor PAC1 was measured using the GloSensor assay^46^ (Promega, Madison, WI). We generated a CHO K1 cell line expressing the respective cAMP-dependent luciferase and the PAC1 receptor. 24 hours before the assay 2500 cells per well were seeded in 100 μl F10 with 10% FBS in white opaque clear-bottom 96-well plates (Costar, Coppell, TX). Before the assay the medium was removed from the wells and replaced with 90 μl per well of Gibco CO_2_-independent medium (Life Technologies) with 10% FBS containing 1% v/v of cAMP GloSensor substrate stock solution (Promega). The cells were incubated in substrate-containing medium at 37 °C in 5% CO_2_ for 2-4 h and equilibrated to room temperature and atmospheric CO_2_ for 30 min. All samples were prepared as duplicates in standard phosphate-buffered saline (PBS) pH 7.4 plus 0.1% (w/v) 3-[(3-cholamidopropyl)dimethylammonio]-1-propanesulfonate (CHAPS) to reduce adsorptive loss of free peptides and 10 μL of 10-fold desired final concentration were added per well. Luminescence was measured for 30 min using a microtiter plate reader and evaluated after the signal reached a plateau (15 min). Dose–response curves were fitted to a four-parameter Hill equation using Prism (GraphPad Software, San Diego, CA). Additional data analysis was performed using proFit (QuantumSoft, Uetikon am See, Switzerland). EC_50_ values are reported as mean and standard deviation of duplicate measurements.

### Preparation and *in vitro* testing of AVATars

All peptides were produced by solid-state synthesis at the MIT Koch Institute Biopolymers and Proteomics Laboratory and purified by high-performance liquid chromatography (HPLC). Biotin (BT) and desthiobiotin (dBT) were incorporated as lysine sidechain-derivatized monomers in the synthesis; polyethylene glycol (PEG)-linked BT was incorporated as a BT-functionalized glutamine monomer spaced using a linker of four PEG units. Identity and purity of each peptide was confirmed by matrix-assisted laser desorption ionization (MALDI)-time of flight mass spectrometry and analytical high-performance liquid chromatography. After lyophilization, the peptides were weighed, dissolved in water to a stock concentration of 1 mM and stored at −80 °C. PAC1 receptor activation by PACAP derived probes was tested with the luminescence assay described above. Activity of PACAP conjugates to BT or dBT were assessed in absence and presence of streptavidin (SA; Thermo Fisher Scientific). For sensor activation studies BT-AVATars were formed by premixing 1 nM PACAP-dBT conjugates with a twofold molar excess of SA and subsequently activated by addition of BT. Reversibility of this sensing was probed by successively adding the following reagents to 1 nM PACAP-dBT: 2 nM SA, 80 nM biotin, 200 nM SA, 8 μM biotin. Further BT-AVATar activation studies were performed using biotinylated BSA (Sigma-Aldrich) containing 100 nM BT or biotinylated cells containing 150 nM BT (preparation see below) as analytes.

To prepare dopamine (DA)-sensitive AVATar, DA was functionalized with a maleimido moiety to provide a facile and thiol-selective conjugation with PACAP substituted at positions 10 and 20 with cysteine residues (PACAP-10,20-Cys). Addition of the maleimido functional group was achieved by reacting DA with N-succinimidyl 3-maleimidopropionate in anhydrous dimethylformamide and further purification using HPLC. The resulting functionalized neurotransmitter, DA-Mal, was further characterized by standard methods (**Supplementary Methods** and **Supplementary Figure 6**). DA-Mal was then reacted with PACAP-10,20-Cys in PBS pH 7.4 using a 6-fold excess of DA-Mal. The resulting conjugate, PACAP-10,20-DA, was purified by HPLC and its molecular structure was confirmed by high-resolution MALDI mass spectrometry. The DA-AVATar was formed by premixing 1 nM PACAP-10,20-DA with 600 nM anti-dopamine IgG antibody (ab6427; Abcam, Cambridge, MA). Dopamine sensitivity of the DA-AVATar was examined using freshly prepared DA (Sigma-Aldrich) stocks in PBS pH 7.4.

### Preparation of biotinylated cell xenografts

HEK293 Freestyle cells were separated into two 10 mL samples 30 million cells/sample. The cells were washed twice with 10 mL ice cold PBS pH 7.4 and resuspended in 1 mL each. The test sample was biotinylated by adding 120 μL of biotinylation reagent (EZ-link NHS-Biotin, Thermo Fisher Scientific). 120 μL of PBS pH 7.4 was added to the control cells. The cells were nutated for 30 minutes at room temperature and another 30 minutes at 4 °C. The cells were washed three times with ice cold PBS pH 7.4 and resuspended in 100 μL artificial cerebrospinal fluid (aCSF). Biotinylation of the cell surface was quantified by labeling biotinylated and control cells with fluorescent SA (SA-DyLight488, Thermo Fisher Scientific). Cells were diluted 1:5 in PBS pH 7.4 and 1/10 of the volume was incubated with 0.1 mg SA-DyLight488 for 30 min at room temperature. Afterwards cells were washed three times with PBS pH 7.4 to remove unreacted SA, diluted 1:10 and DyLight488 fluorescence was quantified using a microtiter plate reader (Excitation: 485 nm, Emission: 538 nm). Cell surface biotinylation was estimated by comparing SA-DyLight fluorescence of cells to a SA-DyLight488 dilution series. All samples were prepared in duplicate. These procedures were performed immediately before implantation experiments described below.

### Animal procedures

All animal procedures were conducted in accordance with National Institutes of Health guidelines and with the approval of the MIT Committee on Animal Care (protocol number 0718-068-21). All experiments were performed with male Sprague-Dawley rats, age 7–9 weeks, supplied by Charles River Laboratories (Wilmington, MA). Seventeen rats were used for *in vivo* imaging experiments described here.

### Preparation of animals for PACAP injection

Seven rats underwent surgery to implant bilateral cannula guides over the caudate putamen region of the striatum (CPu). Anesthesia was induced using 3% isoflurane and maintained using 2% isoflurane with vacuum suction turned on to remove excess anesthetic. The rats were injected subcutaneously with 1.2 mg/kg of sustained release buprenorphine for analgesia. The rats’ eyes were covered with paralube vet ointment (Dechra Veterinary Products, Overland Park, KS) to prevent eyes from drying from exposure to isoflurane. The rats’ heads were shaved and cleaned with alcohol and povidone-iodine prep pads for easy access to the skull. Using sterile surgical equipment, the skin over the skull was retracted and the skull cleaned of tissue so that the sutures on the skull were clearly visible. The holes were drilled through the skull for the cannula guides bilaterally at 3 mm lateral to midline, 0.5 mm anterior from bregma. A small 26 gauge needle was used to puncture the dura in each of the drilled holes, allowing smooth access to the brain parenchyma. The holes were air dried and bilaterally connected custom-made cannula guides (22 gauge PEEK, 6 mm distance between the guides, Plastics One, Roanoke, VA) were inserted into the holes, such that they protruded 1 mm into the brain parenchyma. The guides were then secured to the skull using white dental cement (C&B Metabond, Parkell). Next, a custom head post was attached to the rats’ skulls posterior to the cannula guides implantation site using the white dental cement. After the cement dried, the pink cold cure dental cement (Teets Denture Material, Patterson Dental, Saint Paul, MN) was applied over the white cement to secure the entire implant. Tissue glue was applied to seal the surface connecting the cement and skin areas around the implant. Cannula guides were sealed with dummy cannulas (protruding as far as the guide) to avoid exposure of brain tissue during the recovery period.

### MRI assessment of injected PACAP probes

Seven days after the implantation of the bilateral cannulas, the rats were imaged during PACAP intracranial infusion. For the imaging experiments, animals were anesthetized using 2% isoflurane in oxygen for induction. After numbing the trachea with lidocaine, the animals were intubated intratracheally using the 16 gauge plastic part of a Surflash I.V. catheter (Terumo Medical Products, Somerset, NJ). The rats were then connected to a small animal respirator (Inspira Advanced Safety Ventilator; Harvard Apparatus, Holliston, MA), and fixed via their headposts into a custom-built cradle for imaging with a commercial surface radiofrequency coil for MRI detection (Doty Scientific, Columbia, SC) fitting snugly around the head-post. Breathing rate and end-tidal expired CO_2_ were continuously monitored. Once positioned in the cradle, the anesthesia was lowered to 0.75% isoflurane and the rats were paralyzed with pancuronium (1 mg/kg IP bolus for induction, 2 mg/kg/h IP infusion) to prevent motion artifacts during imaging.

Immediately before each experiment, two injection internal cannulas, designed to protrude 4 mm into the brain past the cannula guides reaching into the CPu (28 gauge PEEK internal cannula, Plastics One, Roanoke, VA, USA), were attached to 25 μL Hamilton glass syringes and prefilled with the appropriate intracranial injection solution (aCSF or 1 μM PACAP in aCSF). Injection cannulas were then lowered, while applying positive pressure of 0.01 μL/min to avoid air bubbles, into the previously implanted bilateral cannula guides. Next, the Hamilton syringes were placed in a remote infuse/withdraw dual syringe pump (PHD 2000 Syringe Pump; Harvard Apparatus). A birdcage resonator coil for radiofrequency excitation in MRI (Doty Scientific) was positioned around the cradle, and the ensemble was inserted into the MRI magnet bore and locked in a position such that the head of the animal was at the center of the bore.

Animals were scanned by MRI using a 7 T 20 cm-horizontal bore scanner (Bruker BioSpin, Billerica, MA) to measure the changes in hemodynamic contrast following intracranial injections. High resolution *T*_2_-weighted anatomical scans of each animal were obtained using a rapid acquisition with relaxation enhancement (RARE) pulse sequence with echo time (*TE*) 44 ms, recycle time (*TR*) 2,500 ms, RARE factor 8, spatial resolution 100 μm × 100 μm × 1 mm, and matrix size 256 × 256 with seven slices. Hemodynamic contrast image series were acquired using a gradient echo planar imaging (EPI) pulse sequence with *TE* 25 ms, *TR* 2,000 ms, spatial resolution 390 μm × 390 μm × 1 mm, and matrix size 64 × 64 with seven slices. Ten minutes of baseline measurement with 4 s per time point were acquired before probe infusion. Following this baseline period, while continuously collecting EPI scans, the infusion pump was remotely turned on to commence intracranial injection of 1 μM PACAP or control solutions at the rate of 0.1 μL/min for ten minutes through the cannulas. EPI scans were collected for an additional 20 minutes after the infusion of the PACAP probe.

MRI data were processed and analyzed using the AFNI software^48^. The AFNI 3dAllineate command was used to align each animal’s EPI data set to the corresponding RARE anatomical image. Each animal’s image data were then aligned to the cannula tip of a reference anatomical MRI. Data from each voxel were normalized to 100, based on the mean of the baseline time period, using AFNI’s 3dcalc command. To identify voxels with significant increases or decreases in BOLD signal, we compared the signal of 10 minutes after the infusion to the baseline in the group concatenated data. For visualization, group signal change maps were overlaid on a reference anatomical image. Time courses were obtained by averaging MRI signal over 1.2 × 1.2 mm regions of interest defined around cannula tip locations in individual animals’ datasets and standard error was calculated across animals using MATLAB. The plateau percent signal change was determined by comparing values observed during the baseline period with values observed after PACAP infusion (minutes 30-40).

### Preparation of animals for imaging biotin-labeled targets

Five rats underwent surgery for implantation of biotinylated and control cells approximately three hours prior to imaging. Craniotomies were performed as described above, except that for these experiments, three holes were drilled through the skull: one for intra-CSF injection of AVATars on the midline just behind the olfactory bulbs (5.2 mm anterior from the bregma), and two for bilateral cell implantations 2 mm lateral to midline and 3.2 mm anterior from bregma. A 1 mm-long guide cannula was implanted at the midline site. To introduce the cells at the lateral sites, two metal cannulas (33 gauge, Plastics One) chosen to minimize damage caused to the brain tissue were attached to 25 μL Hamilton glass syringes and prefilled with the freshly biotinylated or control HEK cells. The injection cannulas were then lowered 2 mm into the brain parenchyma through the two lateral craniotomies and 3 μL of cell suspension (~0.5 million cells) was injected over 30 minutes at 0.1 μL/min. After the cell injection was completed, the metal cannulas were removed, the cell injection holes were dried and sealed, headpost attachment was performed, and a dummy cannula was applied, all as described above.

### *In vivo* imaging with BT-AVATar

Approximately two hours after cell implantation, animals were further prepared for imaging and examined in conjunction with wide-field BT-AVATar infusion. The rats were again intubated, ventilated, and positioned in the MRI cradle with surface coil before being taken off of isoflurane and placed on 0.05 mg/mL dexmedetomidine anesthesia mixed with 1 mg/mL pancuronium for paralysis (1 mL/kg IP bolus for induction, 2 mL/kg/h IP infusion afterwards). An internal cannula designed to protrude 0.25 mm past the preimplanted midline cannula guide was attached to 50 μL Hamilton glass syringe and prefilled with BT-AVATar (2 μM PACAP-10,20,32-dBT mixed with 20 μM SA). The injection cannula was then lowered into the guide, followed by placement of the volume coil over the animal and insertion into the scanner as described above. Animals were scanned by MRI using the same parameters used to validate PACAP itself. Following a ten minute baseline period, infusion of BT-AVATar into the CSF was initiated at a rate of 1 μL/min during continuous scanning.

Data were again processed and analyzed using AFNI. After alignment of each animal’s EPI data to the corresponding anatomical RARE scan, the datasets were all registered in order to align their cell injection sites to those of a reference animal. To eliminate the contribution of nonspecific image fluctuations, we normalized each animal’s time course data to the time course of a reference region in the most posterior slice, assumed not to be affected by AVATar infusion. We then scaled the baseline intensity of each voxel to 100 using AFNI’s 3dcalc command. We excluded from analysis voxels displaying ~50% or greater hypointensity prior to AVATar infusion, which we took to indicate tissue damage in the area of cell injections. To identify voxels with signal changes indicative of AVATar activation, we applied AFNI’s 3dDeconvolve command to identify signal components that displayed a linear increase during AVATar infusion followed by a plateau thereafter. Motion parameters were used as nuisance regressors. For visualization, the resulting beta coefficient maps (in units of percent change) were overlaid on a reference anatomical image. Time courses were obtained by averaging MRI signal from 2.5 × 2.5 mm regions of interest defined around the centroid of cell implantation in individual animals, as judged from anatomical scans. Mean and standard error were calculated across animals using MATLAB. The percent signal change in individual animals was determined by comparing signal values during baseline and during the infusion conditions (minutes 15-30).

### Assessment of contrast agent spread during intra-CSF infusion

Ability of molecular imaging probes to spread broadly through the brain during intra-CSF injection was confirmed using test infusions of gadolinium diethylenetriaminepentaacetic acid (Gd-DTPA, Sigma-Aldrich). In each animal, an intra-CSF cannula guide and headpost were implanted and animals were prepared for imaging as described above, except that the intra-CSF infusion syringe was filled instead with 200 mM Gd-DTPA formulated in aCSF. After insertion of animals into the scanner and acquisition of anatomical scans, *T*_1_-weighted scans suitable for visualizing Gd-DTPA effects were performed using a fast low-angle shot (FLASH) pulse sequence with *TE* 5 ms, *TR* 93.75 ms, spatial resolution 400 μm × 400 μm × 1 mm, and matrix size 64 × 64 with seven sagittal slices. Scans were collected over a ten minute baseline period followed by 30 minutes of contrast agent infusion at a rate of 1 μL/min. The resulting MRI data were processed and analyzed using AFNI. To identify voxels with significant increases or decreases in *T*_1_ signal, we compared the signal during the final ten minutes of Gd-DTPA infusion to the baseline signal from each animal and expressed the result as a percent change in mean intensity. For visualization, a representative signal change map was overlaid on the corresponding sagittal anatomical image.

### Histology

Fluorescent BT-AVATar was prepared by formulating the probes with SA-DyLight 488 (Thermo Fisher Scientific) in place of unlabeled SA. Surgical preparation and biotin-labeled cell implantation were then performed as described above. After infusion procedures identical to those used in the MRI experiments, animals were transcardially perfused with PBS followed by 4% paraformaldehyde in PBS. Brains were extracted, post-fixed overnight at 4 °C, and sectioned the following day. Free-floating sections (50 μm) were cut using a vibratome (Leica VT1200 S, Leica Microsystems GmbH, Wetzlar, Germany), mounted on glass slides with Invitrogen ProLong Gold Antifade Mountant (Fisher Scientific Company, Ottawa, Canada) and protected with a coverslip. BT-AVATar activation in the area of the cell implantation was visualized by the green fluorescence arising from SA-DyLight 488. Independent of biotinylation, the distribution of xenografted cells was also visualized by staining with DAPI (1:1000 dilution from a 1 mg/mL stock, Thermo Fisher Scientific), followed by a wash with PBS plus 0.5% bovine serum albumin (BSA, Sigma-Aldrich). Fluorescence imaging was performed using a confocal microscope (Axio Imager 2; Zeiss, Thornwood, NY).

### Code availability statement

Scripts used for data analysis are available upon reasonable request.

### Data availability statement

Raw MRI datasets generated and/or analyzed for the current study are available from the corresponding author on reasonable request.

## Abbreviations

AVATar: Activatable Vasoprobe for Analyte Targeting
aCSF: Artificial Cerebrospinal Fluid
BT: Biotin
BOLD: Blood Oxygenation Level-Dependent
BSA: Bovine Serum Albumin
CHAPS: 3-[(3-Cholamidopropyl)dimethylammonio]-1-propanesulfonate
CSF: Cerebrospinal Fluid
cAMP: Cyclic Adenosine Monophosphate
dBT: Desthiobiotin
DA: Dopamine
EPI: Echo Planar Imaging
*TE*: Echo Time
Gd-DTPA: Gadolinium Diethylaminetriaminepentaacetic Acid
HPLC: High Performance Liquid Chromatography
MRI: Magnetic Resonance Imaging
MALDI: Matrix-Assisted Laser Desorption Ionization
PBS: Phosphate-Buffered Saline
PACAP: Pituitary Adenylate Cyclase Activating Peptide
PEG: Polyethylene Glycol
RARE: Rapid Acquisition with Relaxation Enhancement
*TR*: Recycle Time
SA: Streptavidin
VSMC: Vascular Smooth Muscle Cell

## ACKNOWLEDGEMENTS

The authors acknowledge grants from the National Institutes of Health (R24 MH109081, U01 NS103470, and UF1 NS107712) to AJ. RO was funded by a fellowship from the Deutsche Forschungsgemeinschaft. AW was funded by the Advanced Multimodal Neuroimaging Training Program at the Massachusetts General Hospital and a fellowship from Harvard-MIT Health Sciences and Technology program. The authors thank Dr. Peter Harvey for help with fluorescence microscopy, and are grateful to Alla Leshinsky of the MIT Koch Institute Biopolymers and Proteomics Laboratory for assistance with peptide synthesis.

## AUTHOR CONTRIBUTIONS

RO performed all *in vitro* experiments. AW performed all *in vivo* experiments. RO, AS, and AJ designed the AVATars. AW, MD, and AJ designed the *in vivo* approaches. AB designed and synthesized tethered dopamine. RO, AW, and AJ wrote the manuscript with assistance from other authors.

## ADDITIONAL INFORMATION

Supplementary information is available in the online version of this paper. Reprints and permissions information are also available online. Correspondence and requests for materials should be addressed to AJ.

## COMPETING FINANCIAL INTERESTS

The authors declare no competing financial interests.

## SUPPLEMENTARY METHODS

### General synthesis-related information

Mal-NHS (**1**) was purchased from Matrix Scientific (Columbia, SC) and used without further purification. All other chemicals were purchased from Sigma-Aldrich. Mass spectra were recorded on Agilent (Santa Clara, CA) 6545 Q-TOF and Bruker (Billerica, MA) Omniflex MALDI-TOF instruments. Nuclear magnetic resonance (NMR) spectra were recorded on a Bruker AVANCE III-400 NMR (400 MHz).

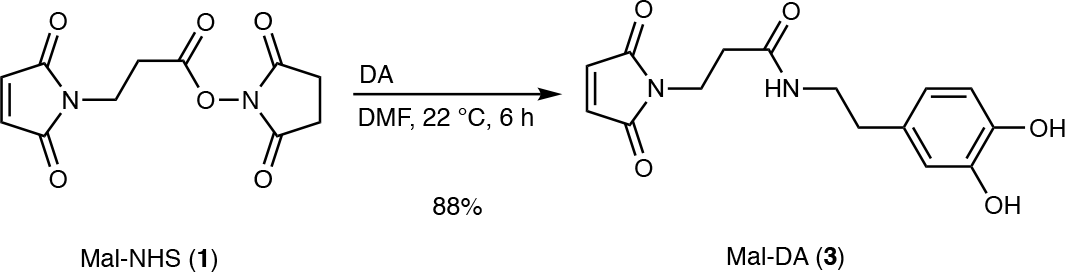

### Synthesis of Mal-DA (3)

Mal-NHS (270 mg, 0.1 mmol) and dopamine hydrochloride (190 mg, 0.1 mmol) were dissolved in anhydrous dimethylformamide (DMF, 2 mL) and after addition of *N*,*N*-diisopropylethylamine (DIEA, 130 mg, 0.1 mmol) the reaction solution was stirred under argon atmosphere at room temperature for 6 h. The resulting solution was treated with deionized water (20 mL) and extracted with EtOAc (3 × 50 mL). The combined organic phases were dried over MgSO_4_ and after removal of all volatiles the resulting residue was purified by high performance liquid chromatography (HPLC, silica-C18, eluent gradient H_2_O:MeCN from 95:5 to 10:90; *t*_R_ = 8.3 min). Yield: 260 mg (88%). MS (HRESI): *m/z* = 303.0991 (theoretical) [M-H]^−^, 303.0928 (experimental) [M-H]^−^. ^1^H NMR (400 MHz, DMSO-*d*_6_) δ 7.97 (t, *J* = 5.8 Hz, 0H), 7.00 (s, 1H), 6.61 (d, *J* = 7.9 Hz, 0H), 6.55 (d, *J* = 1.6 Hz, 0H), 6.47 – 6.30 (m, 0H), 3.59 (t, *J* = 7.3 Hz, 1H), 3.12 (q, *J* = 6.9 Hz, 1H), 2.46 (t, *J* = 7.7 Hz, 1H), 2.30 (t, *J* = 7.3 Hz, 1H). ^13^C NMR (101 MHz, DMSO) δ 170.75, 169.15, 145.03, 143.48, 134.54, 130.15, 119.13, 115.88, 115.44, 40.53, 40.15, 39.99, 39.94, 39.73, 39.52, 39.31, 39.10, 38.89, 34.55, 34.08, 34.03, 0.11.

**Supplementary Table 1.** PAC1 activation by additional PACAP variants.

| Construct | –Log{EC<sub>50</sub>} for PAC1 activation<sup>a</sup> |  |
| --- | --- | --- |
|  | –SA | +SA |
| PACAP | 10.97 ± 0.01 | 10.81 ± 0.01 |
| PACAP-10-BT | 10.20 ± 0.01 | 8.91 ± 0.02 |
| PACAP-10-PEG-BT<sup>b</sup> | 9.45 ± 0.05 | 8.70 ± 0.01 |
| PACAP-20-PEG-BT<sup>b</sup> | 10.7 ± 0.1 | 9.30 ± 0.02 |
| PACAP-10-BT (repurified)<sup>c</sup> | 10.69 ± 0.01 | 9.20 ± 0.04 |

<sup>a</sup> All values in the absence (–SA) or presence (+SA) of streptavidin were measured in duplicate using the PAC1 receptor activation bioassay and presented as mean ± SD.
<sup>b</sup> These variants contain a BT-modified glutamine residue at positions 10 or 20, each with a linker of four polyethylene glycol (PEG) monomers separating the BT moiety from the glutamine sidechain.
<sup>c</sup> PACAP-10-BT was repurified using HPLC to remove potential contaminants.

**Supplementary Figure 1.**
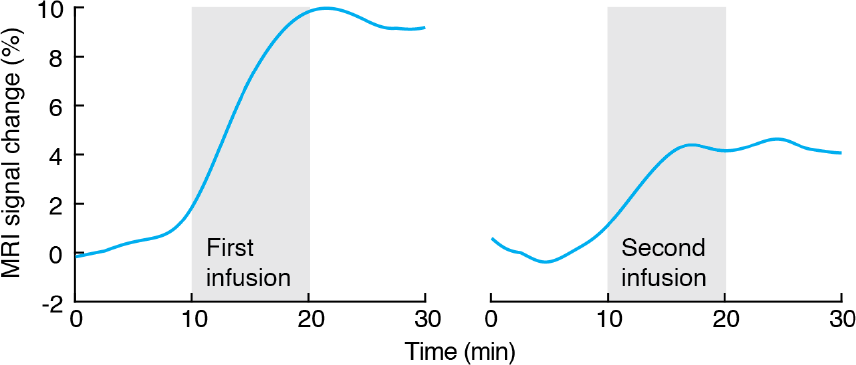
MRI signal changes arising from tandem PACAP injections. Time courses of MRI signal change near the injection region following injections of 1 μM PACAP (0.1 μL/min) over 10 minute intervals within tandem imaging experiments, analogous to main text **Figure 1c-e**. Data presented from a single animal shows responses to PACAP in both experimental periods.

**Supplementary Figure 2.**
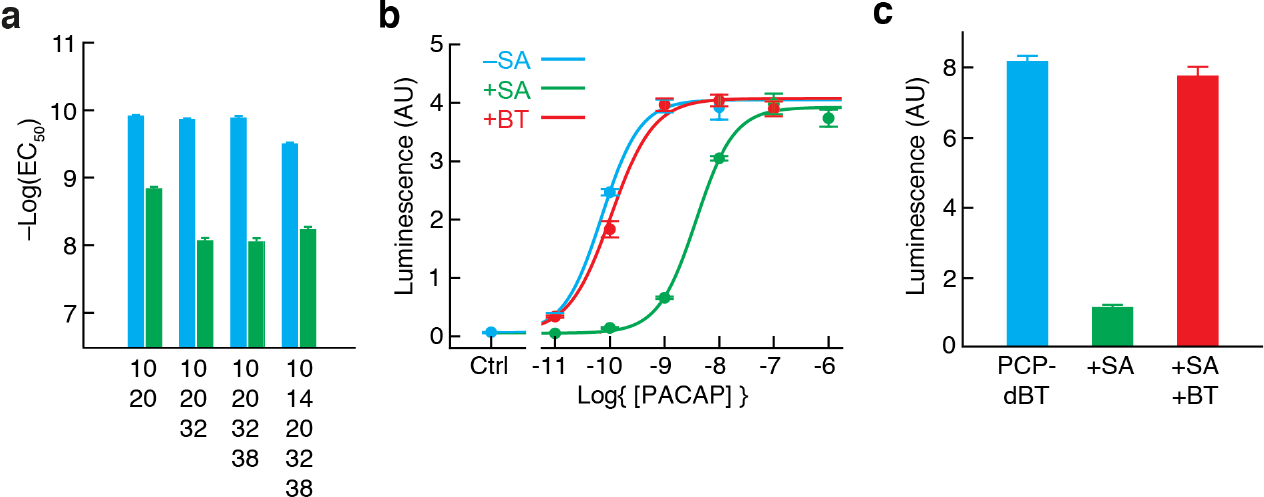
Use of desthiobiotin for BT-AVATar construction. **(a)** EC_50_ values for activation of PAC1 by PACAP derivatives modified with desthiobiotin (dBT) at the residue positions shown, in the absence (cyan) and presence (green) of 200 nM SA. **(b)** Titration of PAC1 receptor activation by PACAP-10,20,32-dBT in the absence (cyan) and presence (green) of 200 nM SA, and in the presence of SA plus 8 μM biotin (BT, red). **(c)** Bioassay output in response to PACAP-10,20,32-dBT with added SA (green) and additionally added BT (red). Error bars denote SD of duplicate measurements.

**Supplementary Figure 3.**
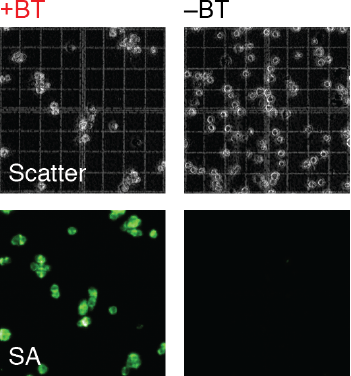
Biotin-labeled and control cells for implantation. Microscopy of biotin-labeled (+BT) and control-treated (−BT) HEK cells by scattered light (top) for visualization of all cells and by SA-Dylight488 (bottom) for visualization of biotin labeling. These cells are equivalent to cells implanted and imaged using BT-AVATar in the experiments of main text **Figure 4**.

**Supplementary Figure 4.**
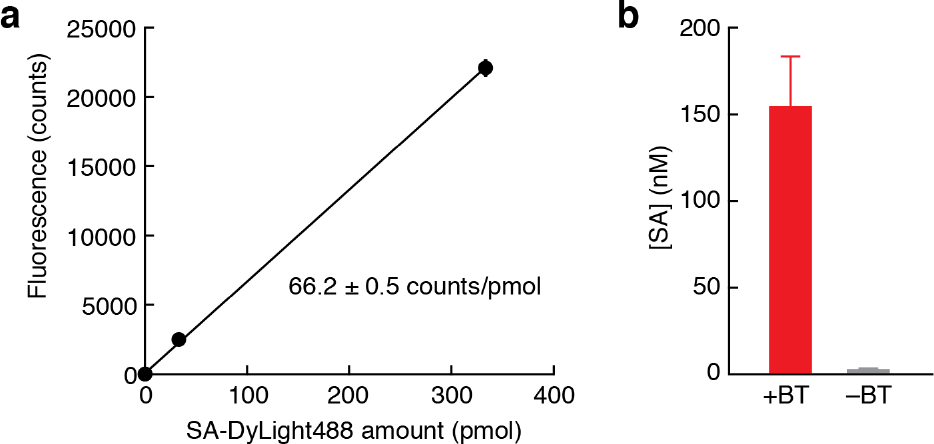
Quantification of streptavidin binding by biotinylated cells. **(a)** Calibration curve showing raw fluorescence output as a function of SA-DyLight488 quantity, yielding the calibration constant indicated. **(b)** Quantification of biotin concentration in cell pellets computed using the calibrated fluorescence assay based on SA-DyLight488 binding to biotin-labeled or control-treated cells.

**Supplementary Figure 5.**
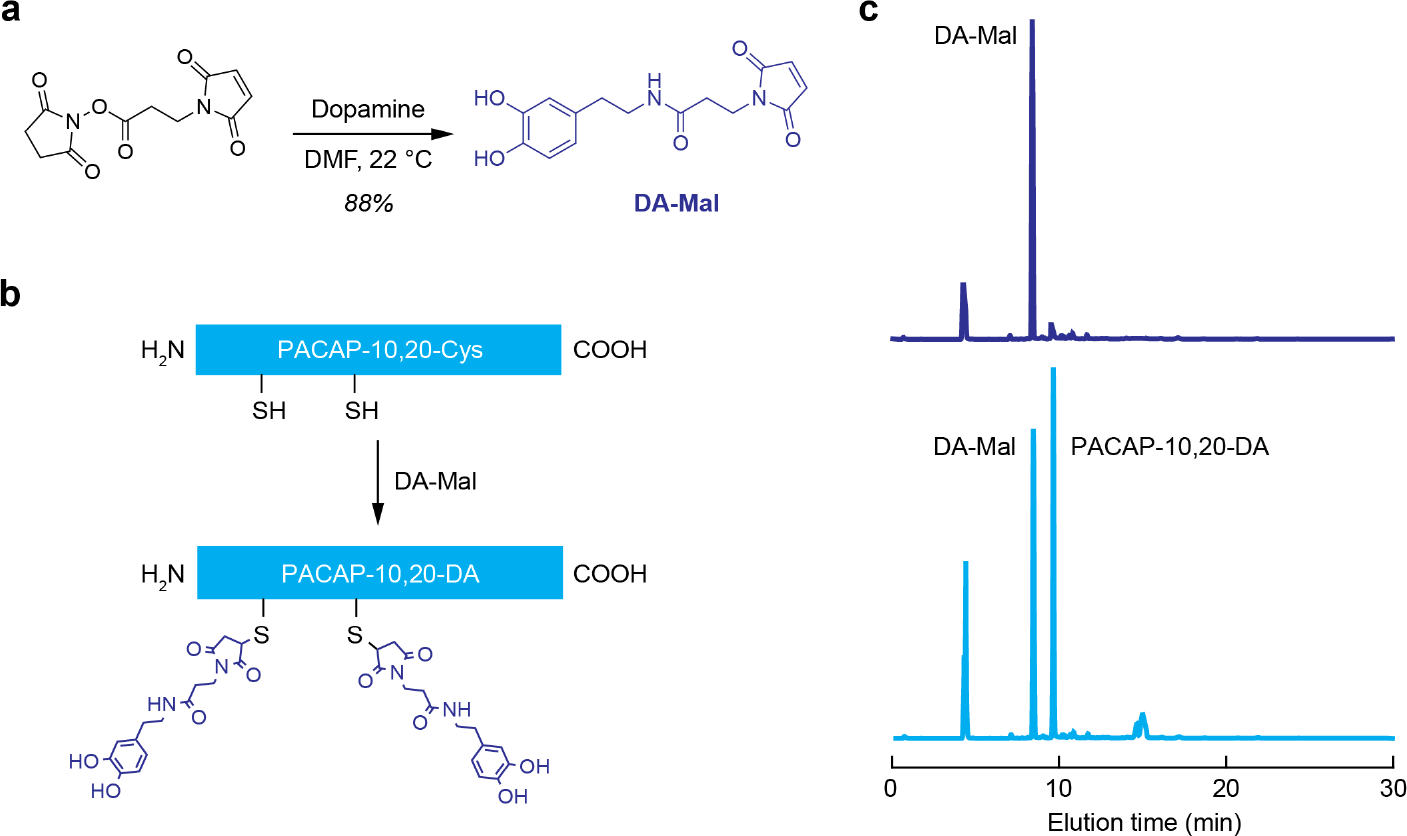
Synthesis of PACAP-10,20-DA. **(a)** Formation of the dopamine-maleimide conjugate DA-Mal by reaction of N-succinimidyl 3-maleimidopropionate (left) with dopamine in DMF. **(b)** The double cysteine mutant of PACAP, PACAP-10,20-Cys was synthesized by solid phase methods and reacted with excess DA-Mal to produce PACAP-10,20-DA as shown. **(c)** HPLC traces showing elution profiles of DA-Mal and PACAP-10,20-DA, following the reaction in **b**.

**Supplementary Figure 6.**
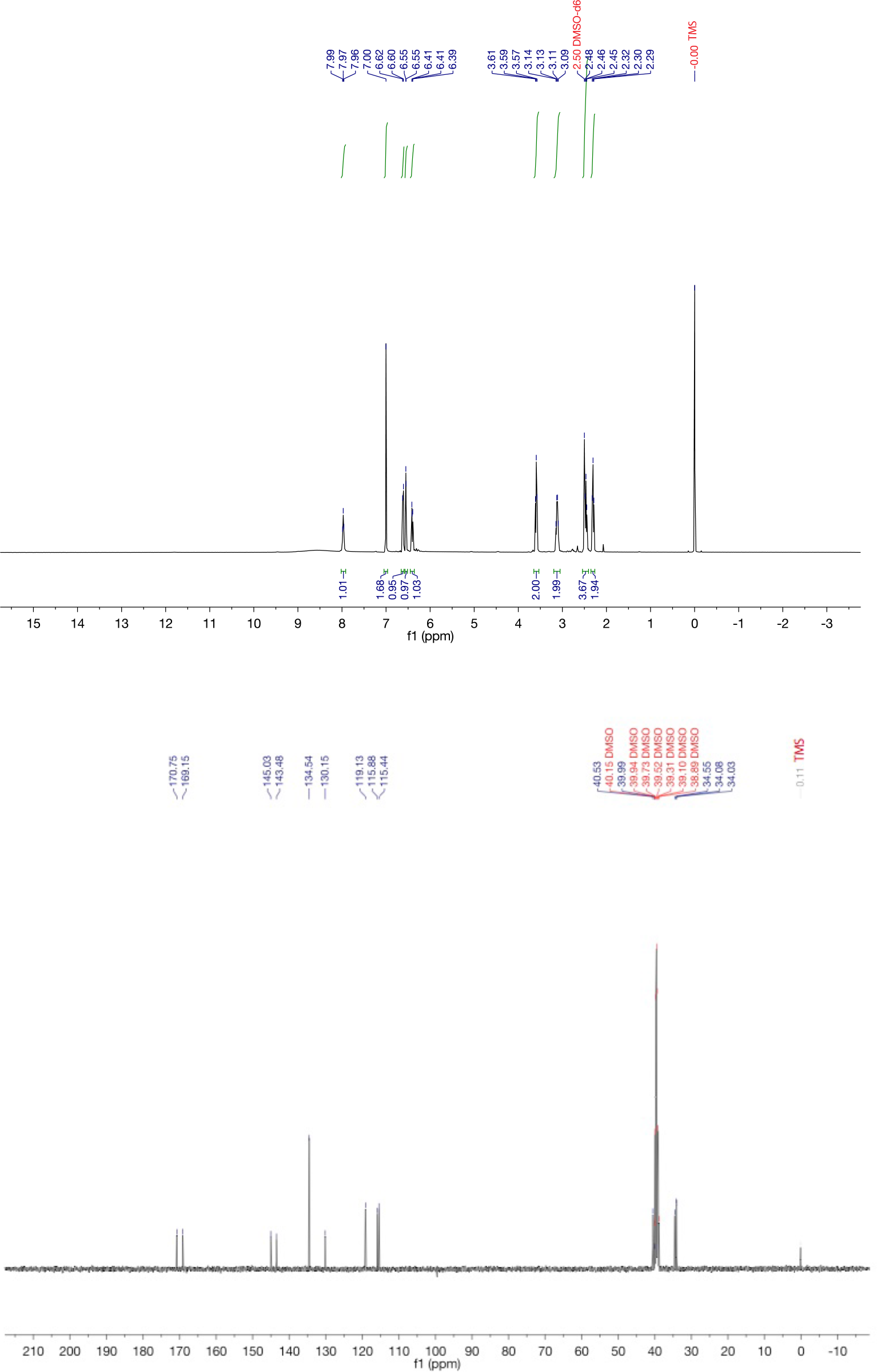
NMR spectra of DA-Mal. Top: ^1^H; bottom: ^13^C.

